# A critical role for Pol II CTD phosphorylation in heterochromatic gene activation

**DOI:** 10.1101/2024.01.26.577526

**Authors:** Amoldeep S. Kainth, Hesheng Zhang, David S. Gross

**Affiliations:** Department of Biochemistry and Molecular Biology Louisiana State University Health Sciences Center Shreveport, LA 71130

**Author notes:** Equal contribution (Amoldeep S. Kainth, Hesheng Zhang). Department of Molecular Genetics & Cell Biology University of Chicago Chicago, IL 60637.

## Abstract

How gene activation works in heterochromatin, and how the mechanism might differ from the one used in euchromatin, has been largely unexplored. Previous work has shown that in *SIR*-regulated heterochromatin of *Saccharomyces cerevisiae*, gene activation occurs in the absence of covalent histone modifications and other alterations of chromatin commonly associated with transcription. Here we demonstrate that such activation occurs in a substantial fraction of cells, and this raises the possibility that an alternative activation pathway might be used. We address one such possibility, Pol II CTD phosphorylation, and explore this idea using a natural telomere-linked gene, *YFR057w*, as a model. Unlike covalent histone modifications, CTD phosphorylation at Ser2, Ser5 and Ser7 is abundant at the drug-induced heterochromatic gene. Particularly enriched relative to the euchromatic state is Ser2 phosphorylation. Consistent with a functional role for Ser2P, *YFR057w* is negligibly activated in cells deficient in the Ser2 CTD kinases Ctk1 and Bur1 even though the gene is strongly stimulated in *ctk1Δ bur1* cells when it is placed in a euchromatic context. Collectively, our results are consistent with a critical role for Ser2 CTD phosphorylation in driving Pol II recruitment and transcription of a natural heterochromatic gene – an activity that may help supplant the need for histone epigenetic modifications.

## Introduction

In eukaryotes, chromatin provides the center stage to orchestrate the complex symphony of transcription. Classic models subdivide chromatin into two types – an open, accessible state called euchromatin that is conducive to transcription and a cytologically condensed state called heterochromatin that is largely reticent to transcription (Gross et al., 2015). It has long been accepted that underlying the function of these two domains are disparate sets of covalent histone modifications that constitute the so-called ‘histone code’ (Strahl and Allis, 2000; Jenuwein and Allis, 2001; Bannister and Kouzarides, 2011). Euchromatin is enriched in activating marks such as methylation of H3K4, H3K36 and H3K79 and acetylation of H3K9, H3K27 and H4K16 as well as other lysine residues on H2A, H2B, H3 and H4. These modifications facilitate the binding and activity of transcription factors, RNA polymerase and other regulatory proteins that drive gene transcription (reviewed in Smolle and Workman, 2013; Rothbart and Strahl, 2014; Lawrence et al., 2016). In contrast, heterochromatin is impoverished in H3 and H4 acetylation while enriched in repressive marks such as H3K9, H3K27 and H4K20 methylation. The latter serve as docking sites for proteins such as HP1 and the PRC2 Polycomb complex that compact chromatin and suppress transcription (reviewed in Grewal and Moazed, 2003; Gross, 2015; Gross et al., 2015).

Several studies have questioned the role of these canonical histone modifications. Activation of heterochromatic genes in the budding yeast *Saccharomyces cerevisiae* occurs in the absence of many classical epigenetic marks (Sekinger and Gross, 2001; Zhang et al., 2014). Likewise, in *Arabidopsis*, stress-induced activation of heterochromatic transgenes occurs without reversal of repressive marks (Tittel-Elmer et al., 2010). In addition, developmentally regulated genes in *Drosophila* can be transcribed in the absence of H3K4me3 and H3K36me3 (Perez-Lluch et al., 2015) while essentially normal transcriptional regulation in that organism can occur in the complete absence of H3K4 methylation (Hodl and Basler, 2012). In mouse, major satellite repeat genes are transcriptionally induced in response to proteasome inhibition in the absence of changes in either repressing or activating marks (Natisvili et al., 2016). Activation of heterochromatic transcription, at least in certain cases, can be attributed to the co-occupancy of gene-specific transcription factors and pioneer factors with repressive chromatin (Sekinger and Gross, 1999; Sekinger and Gross, 2001; Chen and Widom, 2005; Perez-Lluch et al., 2015; Bondra and Rine, 2023). Therefore, instead of being the defining feature of transcriptional changes, covalent histone modifications work in conjugation with other modes of regulation. However, as these studies employed population-wide assays, the prevalence and frequency of transcription changes in individual cells independent of histone covalent modification remains unknown.

The notion of heterochromatin as being structurally monolithic is also contested by recent findings that both constitutive and facultative types of heterochromatin (HP1- and PRC2-containing, respectively) are formed by liquid-liquid phase separation. As such, these heterochromatic domains form membraneless compartments that exhibit high internal diffusion and selective permeability of ligands (Larson et al., 2017; Strom et al., 2017; Markaki et al., 2021). Congruent with its structural pliability, heterochromatin is also functionally dynamic. As an example, *SIR*-mediated silencing in yeast is inversely correlated to the strength the gene-specific transcription factors that occupy it (Wang et al., 2015). Moreover, genes residing in heterochromatic domains in higher eukaryotes respond to diverse cellular signals and environmental cues including stress, development, aging and disease progression (Tittel-Elmer et al., 2010; Ochoa Thomas et al., 2020; Okabe et al., 2020; Wasserzug-Pash et al., 2022; Zhang et al., 2022; Anderson et al., 2023). In contrast to the conventional notion of heterochromatin being a fixed and inert structure, these studies reveal the structural and functional plasticity of heterochromatin that allows it to respond dynamically to diverse stimuli. However, how exactly transcription is achieved in heterochromatin and the uniformity of activation across the cell population remains unknown.

A second transcriptional regulatory system involves covalent modification of the C-terminal domain (CTD) of RNA Pol II. Like the N-terminal tails of the core histones, the Pol II CTD is intrinsically disordered; it is comprised of heptad repeats with a consensus sequence of YSTPSPS. Human Pol II contains 52 heptad repeats (the distal half of which are degenerate) while budding yeast contains 26 (almost all of which match the consensus) (Harlen and Churchman, 2017). Reversible phosphorylation of the CTD plays a key role in regulating different phases of the transcription cycle as well as in regulating histone covalent modifications that occur during Pol II elongation (Harlen and Churchman, 2017). Phosphorylation of Ser5, mediated by Cdk7/Kin28, triggers transcription initiation, Pol II promoter escape and early elongation. In higher eukaryotes, Ser5-phosphorylated Pol II typically pauses 20-50 bp following early elongation. After receipt of a second signal, often under developmental or environmental control, Cdk9/P-TEFb-mediated phosphorylation of Ser2 triggers release of Pol II from its paused state into productive elongation (Buratowski, 2009; Fuda et al., 2009). Phosphorylation of Ser7, principally catalyzed by Cdk7/Kin28, has been linked to the regulation of genes encoding small nuclear RNAs (reviewed in Egloff, 2012).

Similar to covalent histone modifications, CTD modifications are proposed to create an interacting moiety for gene regulatory factors. In this case, the molecules recruited include elongation factors, chromatin remodelers, mRNA capping enzymes, splicing factors, and cleavage and polyadenylation factors (Saunders et al., 2006; Govind et al., 2010; Tietjen et al., 2010; Harlen and Churchman, 2017). CTD phosphorylation has also been linked to the formation of phase-separated transcriptional condensates specifically associated with enhancers and promoters (Boehning et al., 2018) or splicing complexes (Guo et al., 2019). Although the role of CTD modifications in gene activation of euchromatin has been studied extensively, little is known about their impact on gene activation in heterochromatin. Here we explore the role of Pol II CTD phosphorylation in fostering transcription of heterochromatin in budding yeast, a context in which histone post-translational modifications (PTMs) play a minimal role.

## Results

### The model system: transgenic and natural yeast genes under *SIR* regulation

In previous work, we have shown that heterochromatic yeast genes under the regulation of the Sir2/3/4 complex, both naturally occurring and transgenic, are strongly repressed relative to their euchromatic counterparts (as much as 100-fold) (Sekinger and Gross, 1999; Sekinger and Gross, 2001; Zhang et al., 2014). Nonetheless, they are strongly inducible. For example, mRNA levels of *YFR057w*, a natural subtelomeric gene located on the right arm of chromosome VI, are several hundred-fold induced following exposure of cells to the drug cycloheximide (CX). Likewise, those of *hsp82-2001* – an allele of *HSP82* flanked by integrated *HMRE* transcriptional silencers (Figure 1A) – are several hundred-fold induced following exposure of cells to acute thermal stress (Sekinger and Gross, 1999; Zhang et al., 2014). Notably, induction of both genes occurs in the absence of chromatin alterations commonly linked to transcriptional activation, including nucleosome eviction, histone acetylation, methylation of H3K4, H3K36 and H3K79 residues, and replacement of H2A with the H2A.Z variant Htz1 (schematically summarized in Figure 1B).

**Figure 1.**
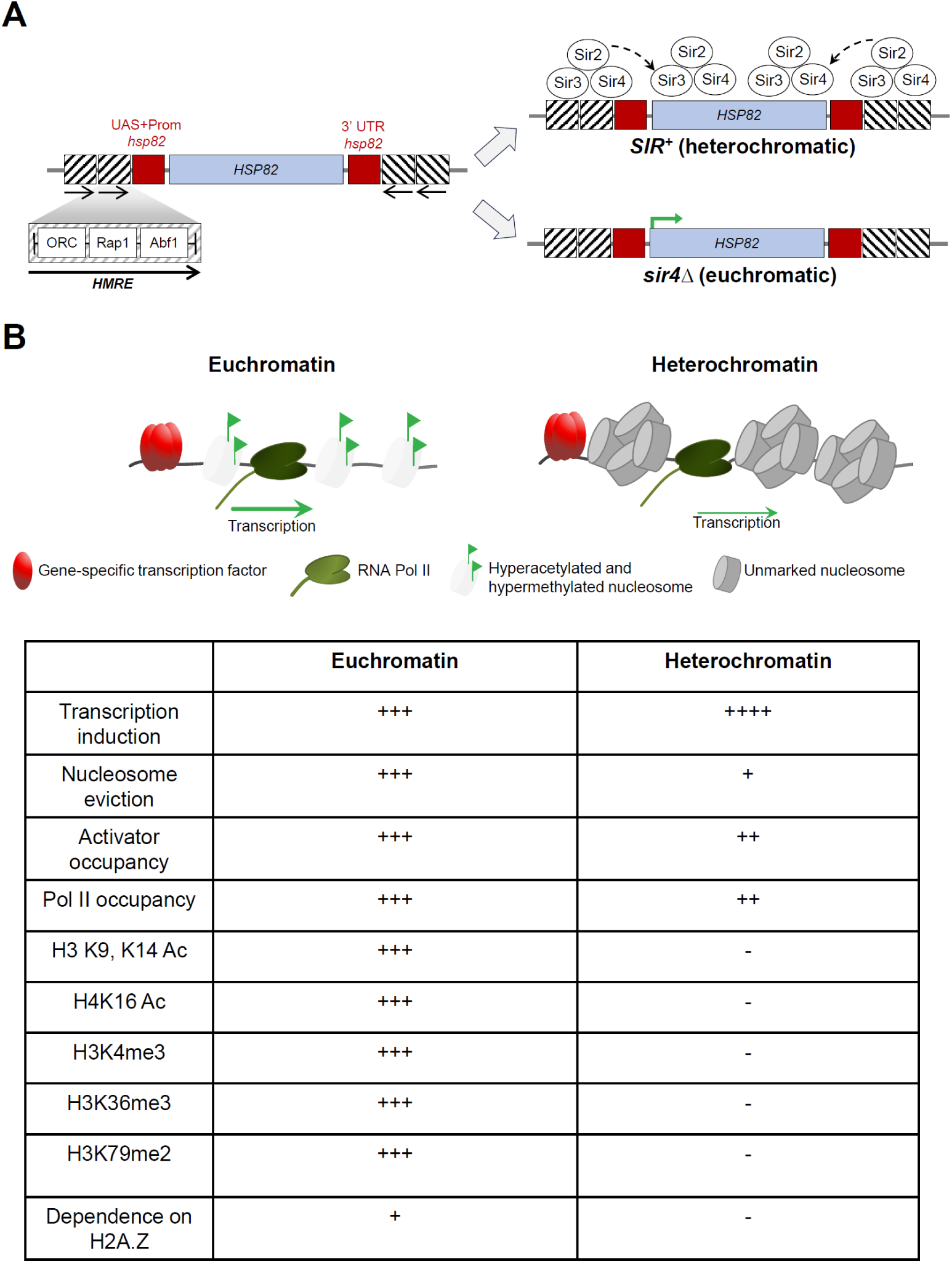
Heterochromatic yeast genes are activated in the absence of chromatin alterations commonly associated with transcription. **A.** Physical map of the model *hsp82-2001* transgene, an *HSP82* allele flanked by tandem *HMRE* silencers (144 bp sequence elements containing binding sites for the indicated factors separated by 6 bp spacers) targeted 675 bp upstream of the transcription start site and 150 bp downstream of the ORF in the orientations indicated (Sekinger and Gross, 1999). Also shown is a schematic representation of the recruited Sir2/3/4 complex observed at this chromosomal locus in *SIR^+^*, but not *sir41*, cells (Sekinger and Gross, 2001; Zhang et al., 2014). **B.** Schematic model and summary of key parameters associated with the transcriptional activation of model euchromatic and heterochromatic yeast genes (Sekinger and Gross, 1999; Sekinger and Gross, 2001; Zhang et al., 2014). Note that ‘transcription induction’, ‘nucleosome eviction’ and ‘dependence on H2A.Z’ refer to fold-change; all other entries refer to absolute levels.

### Widespread expression of the heterochromatic *hsp82* heat shock gene takes place following exposure of cells to acute thermal stress

The above observations suggest that heterochromatic gene activation in *S. cerevisiae* can occur in the context of high nucleosome density and stability as well as in the absence of detectable covalent histone modifications. However, these findings may simply reflect the fact that a very small fraction of the cell population undergoes transcriptional activation and that changes in histone modification state linked to rare activation events are undetectable. To rule this out, we engineered yeast to express eGFP under the control of the heat-shock inducible *hsp82-2001* promoter in both *sir4Δ* and *SIR^+^* contexts to permit evaluation of transgene expression in single cells using fluorescence microscopy. As above, *hsp82-2001-eGFP* is under *SIR* regulation due to the presence of integrated, flanking *HMRE* silencers (schematically illustrated in Figure 2A). Deletion of *SIR4* conferred a mild growth defect at 30°C that was suppressed at 37°C (Figure 2B), consistent with previous observations (https://www.yeastgenome.org/). Due to the basal transcription driven by this promoter (McDaniel et al., 1989), nearly all cells were eGFP-positive in the euchromatic (*sir41*) context (Figure 2C). By contrast, virtually no fluorescent cells were observed in the *SIR^+^* context, confirming that gene silencing efficiently occurs at this reporter.

**Figure 2.**
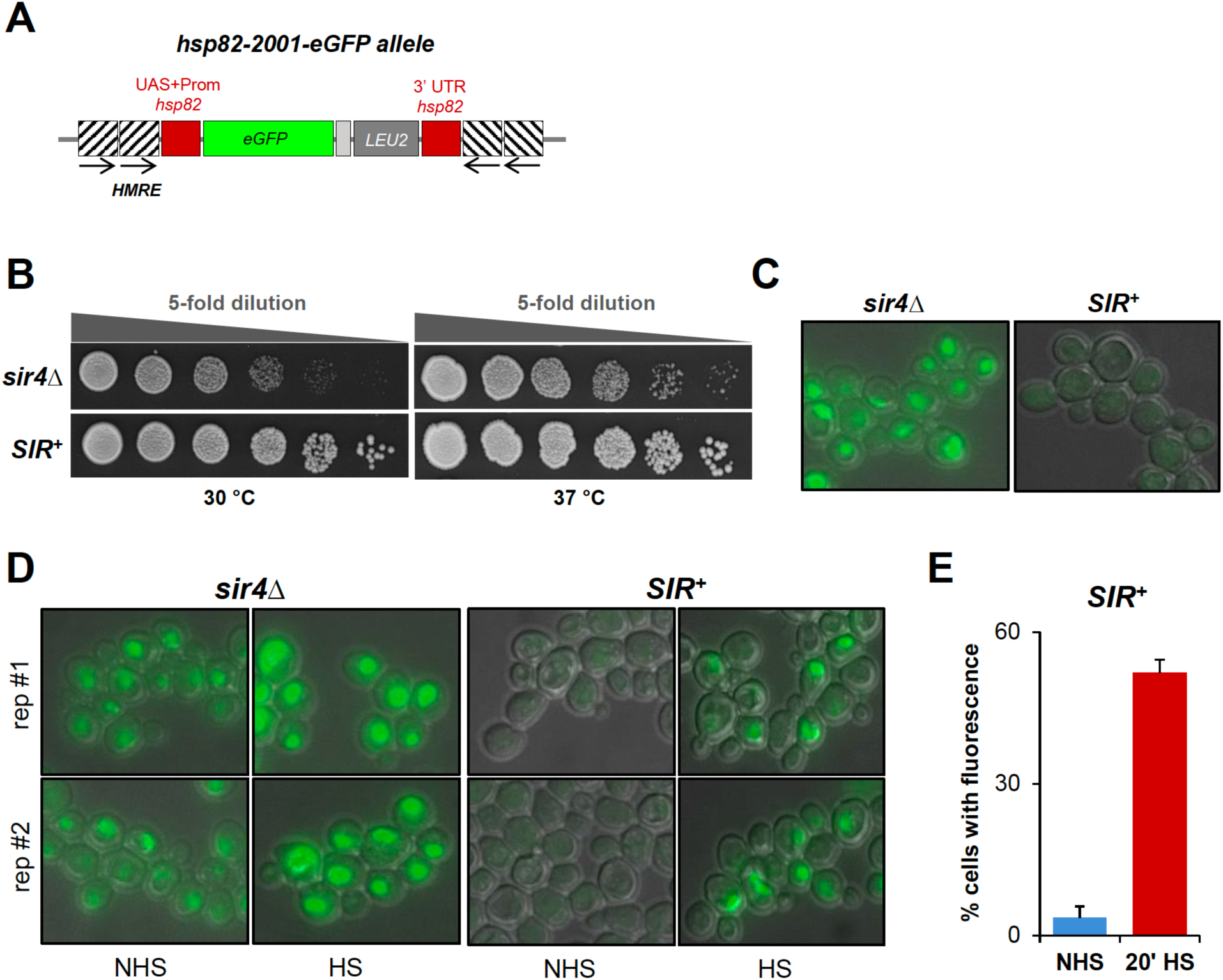
Fluorescence microscopy reveals that both euchromatic and heterochromatic alleles of *hsp82* undergo activation in cells exposed to acute thermal stress. **A.** Map of the *hsp82-2001-eGFP* promoter-gene fusion. *HMRE* silencers are symbolized as in Figure 1. In this construction, *eGFP-LEU2* was integrated at the *hsp82-2001* locus, replacing the gene’s ORF. **B.** Spot dilution growth assay of cells bearing *hsp82-2001-eGFP* in euchromatic (*sir4Δ*) and heterochromatic (*SIR^+^*) contexts grown at 30°C and 37°C. **C.** Micrographs of cells bearing *hsp82-2001-eGFP* in euchromatic and heterochromatic contexts under control, non-heat shock conditions. **D.** Micrographs of cells bearing *hsp82-2001-eGFP* in euchromatic and heterochromatic contexts under the indicated conditions. NHS, non-heat shock; HS, heat shock. **E.** Percentage of *SIR^+^* cells bearing *hsp82-2001-eGFP* with detectable GFP fluorescence under the indicated conditions.

Next, we tested the effect of inducing *hsp82-2001-eGFP* by thermal stimulus. As observed above, the euchromatic *hsp82-2001-eGFP* gene was uniformly expressed across the cell population under non-heat-shock (NHS; 30°C) conditions (Figure 2D). Following 20 min exposure to heat shock (HS; 39°C), GFP expression was considerably higher (compare NHS vs. HS images in *left* panels). In contrast to this uniform expression, expression of the *SIR*-silenced *hsp82-2001-eGFP* gene was strongly variegated under NHS conditions, as it was detectable in only 3% of cells. This is consistent with the incidental escape from *SIR* silencing reported previously (Dodson and Rine, 2015). Importantly, the proportion of cells expressing eGFP increased to 50% following a 20 min heat shock (Figures 2D and 2E) and to 65% following 60 min heat shock (data not shown).

Due to basal expression of *hsp82-2001-eGFP* in the euchromatic context, microscopy analyses are unable to discern the heat shock-induced increase in the proportion of GFP-positive cells in the population. Microscopy is also limiting since it permits analysis of a comparatively small number of cells. To overcome these limitations, we performed a flow cytometry analysis of both *SIR*^+^ and *sir4Δ* cells (see Figure 3A for experimental strategy). As illustrated in Figure 3B, there is a similar distribution of cells in the forward scatter channel (FSC-A) across the six experimental conditions, indicating a uniform distribution of cell size and absence of clumping. The two-dimensional spread plots also depict GFP-positive events at the single cell level. While we observed a largely homogenous activation of eGFP in the euchromatic *sir4Δ* context for cells exposed to either 20- or 60-min heat shock, heat shock-induced activation in the heterochromatic *SIR*^+^ context resulted in at least two discrete subpopulations (Figures 3B and 3C). Moreover, a similar fraction of cells exhibited an increase in eGFP expression in response to heat shock in both *sir4Δ* and *SIR*^+^ contexts as revealed by Overton histogram subtraction analysis (Figure 3D), consistent with the above microscopy results. Therefore, the absence of detectable covalent histone modifications at the heterochromatic *hsp82* transgene cannot be attributed to the fact that only a miniscule fraction of the cell population expresses this gene. Instead, the results shown in Figures 2 and 3 are consistent with ≥50% of cells capable of expressing this gene under the inducing condition, supporting the claim that heterochromatic gene activation can occur in the absence of canonical histone marks.

**Figure 3.**
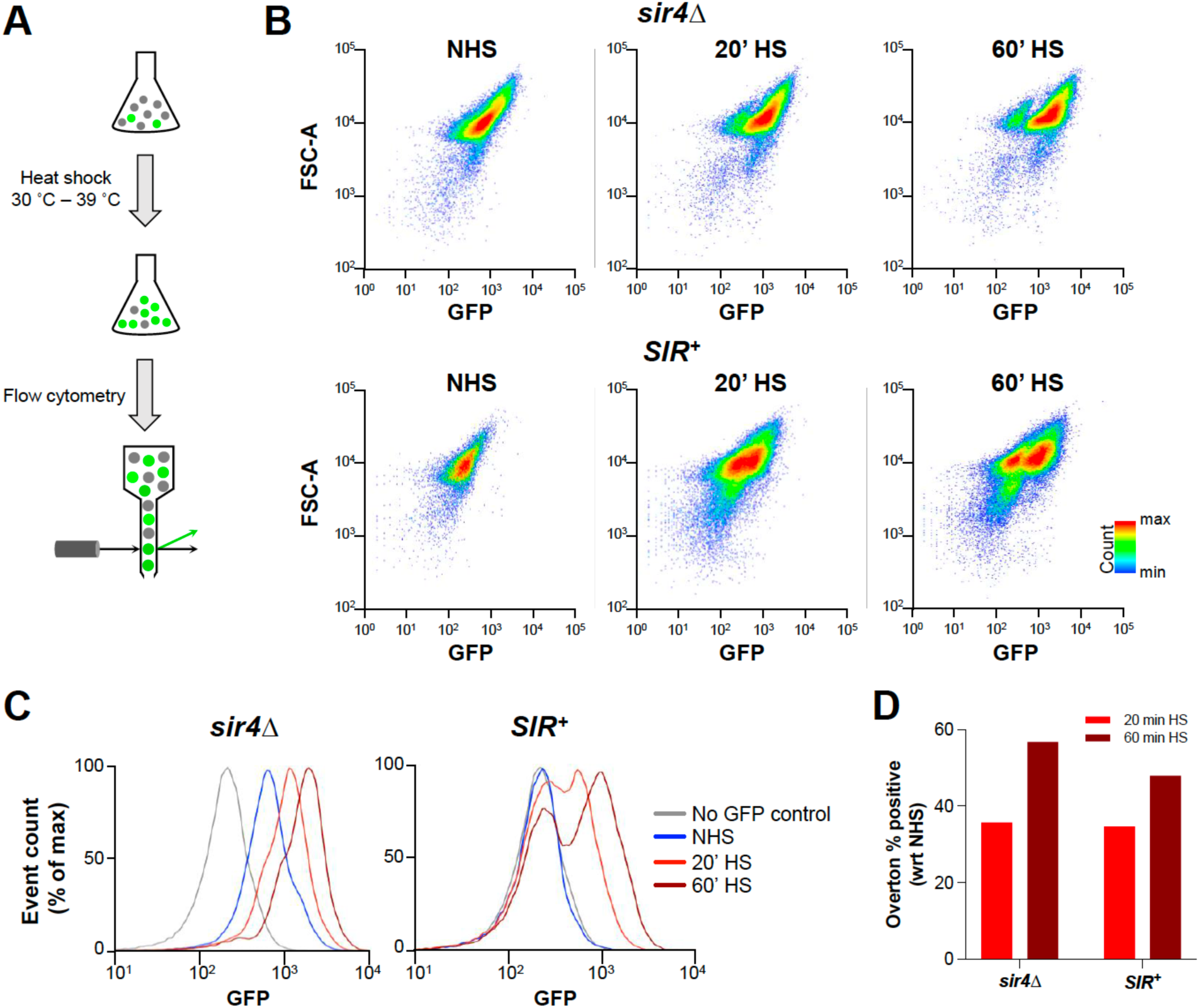
Heterochromatic *hsp82-2001-eGFP* expression is induced in a substantial fraction of cells following exposure to heat shock. **A.** Experimental setup used for flow cytometry analysis to evaluate the heat-shock induced changes in GFP expression in the population of cells. **B.** 2D scatter plot of flow cytometry events on the forward scatter (FCS-A) and GFP channel of the population of cells (comprised of ∼50,000 cells each) bearing *hsp82-2001-eGFP* in both euchromatic (*sir4Δ*) and heterochromatic (*SIR^+^*) contexts under the indicated conditions. The color bar indicates the density of event counts from low (blue) to high (red). **C.** Histograms depicting the frequency distribution as a function of GFP fluorescence intensity of *hsp82-2001-eGFP*-containing cells subjected to the indicated conditions. A yeast strain without any GFP was used to set the baseline. **D.** Overton histogram subtraction analysis of the heat shocked *sir4Δ* and *SIR^+^* cell populations with respect to the non-heat shocked cells. The results indicate an approximate 50-60% increase in the fraction of cells with enhanced GFP expression following either a 20-min or 60-min HS.

### Heterochromatic gene activation is correlated with heightened Pol II CTD phosphorylation

The above observations in combination with previous ones raise the possibility that a gene may use a different pathway for its transcriptional activation depending on its chromatin environment. To test this idea, we investigated the phosphorylation state of the Pol II CTD in the context of heterochromatin. Like the histone N-terminal tails, the Pol II CTD is highly conserved and is subject to reversible covalent modification. Also like the histone tails, it is intrinsically disordered and can serve as a flexible scaffold that allows for many different interactions with target proteins (reviewed in Harlen and Churchman, 2017). While CTD phosphorylation has been demonstrated to participate in histone modifications (reviewed in Tomson and Arndt, 2013), it is possible that other mechanisms, including CTD phosphorylation-triggered recruitment of elongation factors (Zhang et al., 2012; Harlen and Churchman, 2017), may foster chromatin transcription independently of covalent histone modification.

To provide the strongest possible model for this test, we investigated the CTD phosphorylation state of Pol II at the naturally occurring telomere linked *YFR057w* gene which, as mentioned above, has been shown to be strongly repressed by the Sir2/3/4 complex yet potently induced following exposure to cycloheximide (Zhang et al., 2014). This induction is consistent with the presence of a consensus sequence for Stb5 (forms a heterodimer with the pleiotropic drug activator Pdr1) upstream of the gene. Ser5P phosphorylation of the CTD, unlike covalent histone modification, was readily detectable at the subtelomeric gene, both prior to and following addition of the drug (Figure 4, A and B, black bars). In fact, Ser5-phosphorylation – abundant at the 5’-end of protein-encoding genes and linked to early elongation events – was present at comparable levels in heterochromatic (*SIR^+^*) and euchromatic (*sir2Δ*) contexts. Likewise, Pol II was Ser7-phosphorylated within both promoter and coding regions in heterochromatin, resembling and perhaps even exceeding its level of modification in euchromatin (Figure 4, C and D).

**Figure 4.**
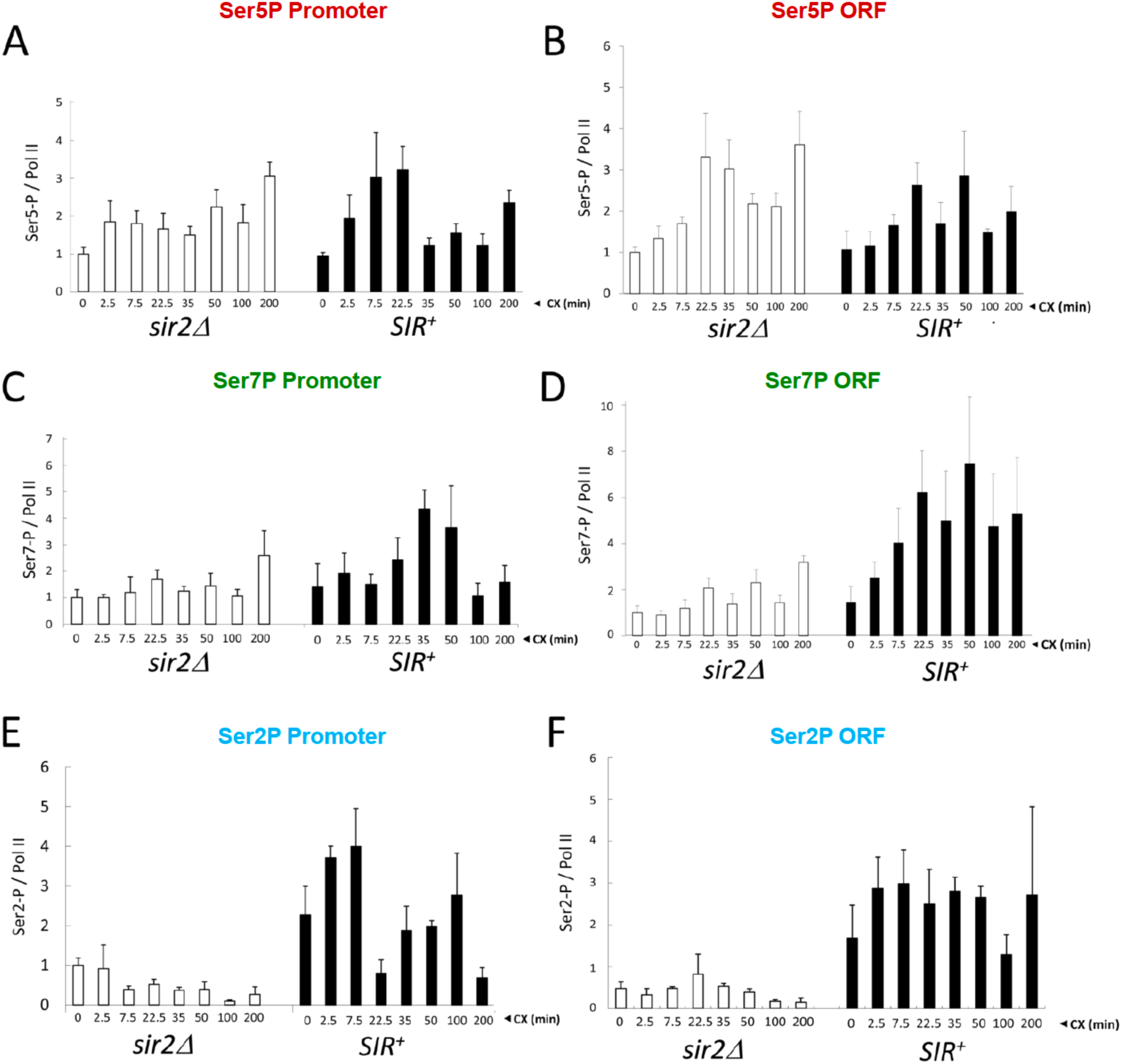
The Pol II CTD is heavily phosphorylated at the transcriptionally activated, subtelomeric *YFR057w* gene. ChIP-qPCR analysis of the Ser5-, Ser7- and Ser2-phosphorylated isoforms of Pol II within the *YFR057w* promoter (panels A, C, E) and coding region (B, D, F) in isogenic *SIR^+^* and *sir2Δ* and cells (strains BY4741 and LG2883, respectively) at the indicated times following addition of 200 μg/ml cycloheximide (CX). Depicted are normalized PhosphoCTD/Rpb1 quotients as means + S.D. (N=2; qPCR=4).

Most intriguing, Ser2-phosphorylated Pol II, linked to the productively elongating form of polymerase, was substantially enriched at the heterochromatic gene relative to its euchromatic counterpart (*P*≤0.005 at most time points [two-tailed *t* test]; Figure 4, E and F). Enrichment of the Ser2P form of Pol II may be particularly important in heterochromatic *YFR057w* activation given that Pol II is only minimally modified at this residue in the euchromatic gene. The latter is consistent with a genome-wide analysis of actively transcribed genes showing that short genes accumulate significantly lower levels of Ser2P Pol II than do long ones (Bataille et al., 2012). (The *YFR057w* ORF is 456 bp.) Thus, both CTD phosphorylation and the previously observed histone modification profiles of *YFR057w* in *sir2Δ* cells – high levels of H3 and H4 acetylation, H3 K4, K36 and K79 methylation, and Htz1 (Zhang et al., 2014) – are typical for an actively transcribed euchromatic gene.

### Ser2 CTD kinase activity is essential for heterochromatic gene activation

As we observed a higher level of Pol II CTD phosphorylation, especially Ser2P, in *SIR^+^* vs. *sir2Δ* cells, we next tested if heterochromatic gene activation has an enhanced requirement for this modification compared to euchromatin. We compared *YFR057w* mRNA levels in WT cells with those in *ctk1Δ* mutants lacking the principal Ser2 CTD kinase, Ctk1. As shown in Figure 5A, loss of bulk Ser2 CTD kinase activity virtually abolished heterochromatic *YFR057w* transcript levels in cells exposed to the drug (*SIR^+^*panel). The *ctk1Δ* mutation also diminished expression of the euchromatic *YFR057w* gene (*sir2Δ* panel), but its effect was less severe (∼3-fold) and may in part arise from 3’- end defects that destabilize the transcript (Ahn et al., 2004). The greater fold-repression of *YFR057w* RNA levels elicited by loss of Ctk1 in *SIR^+^ vs. sir2Δ* cells (Figure 5A, right panel) is therefore consistent with a heterochromatic-specific requirement for Ser2 CTD phosphorylation.

**Figure 5.**
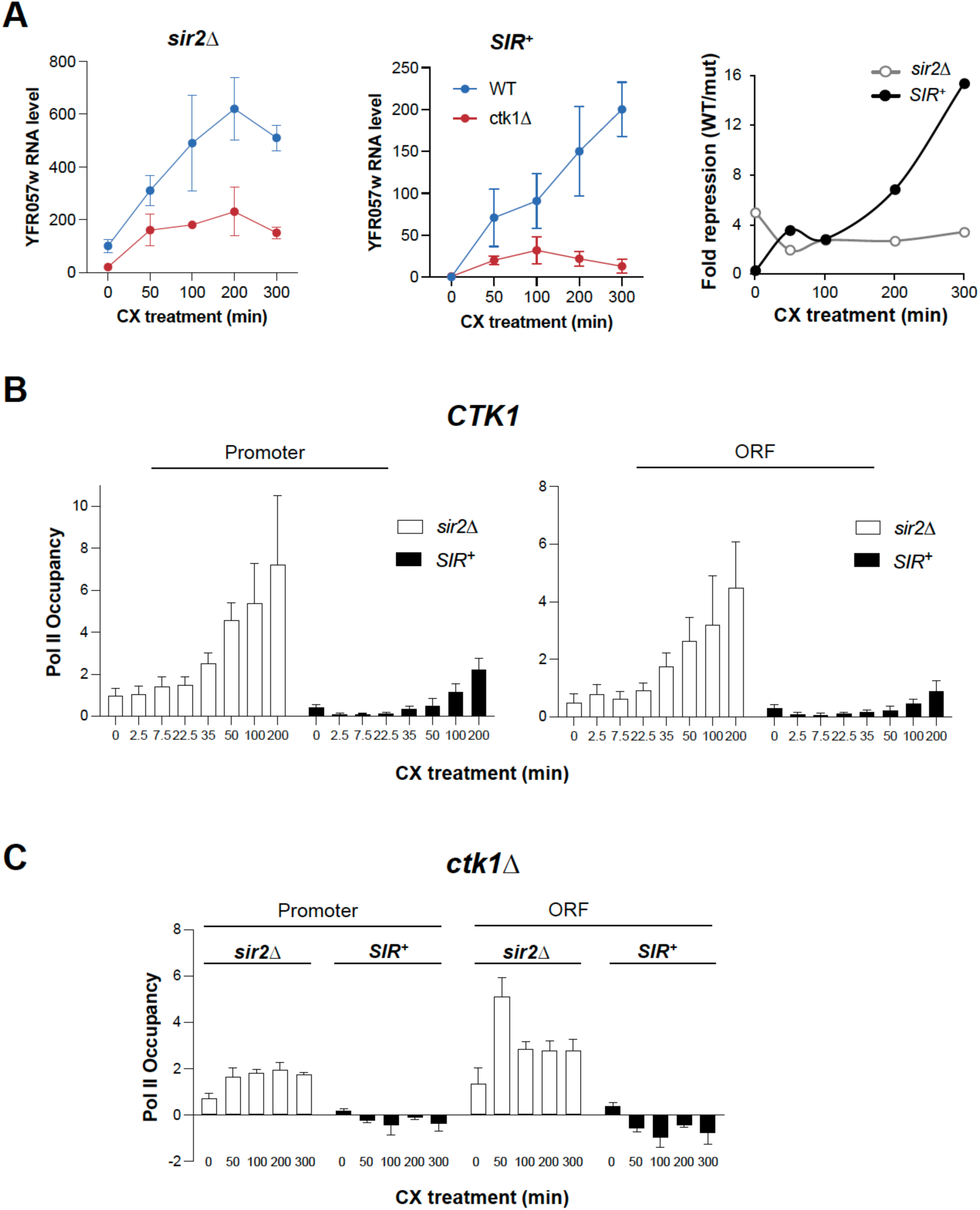
Heterochromatic gene activation is strongly suppressed in *ctk1Δ* cells. **A.** Expression assays of *YFR057w* in isogenic *sir2Δ* and *SIR^+^* cells bearing either *CTK1* or *ctk1Δ* as indicated (strains HZY103, HQY1223, HZY104, and HQY1190, respectively). The left two panels depict an RT-qPCR analysis; shown are means + S.D. (N= 3; qPCR=6). Right panel: fold reduction in *YFR057w* RNA levels arising from the *ctk1* deletion. **B.** ChIP-qPCR analysis of Pol II occupancy within the *YFR057w* promoter and ORF in *sir2Δ* and *SIR^+^* cells exposed to 200 μg/ml cycloheximide for the indicated times. Occupancy of the non-induced promoter (*sir2Δ*) was set to 1.0; all other occupancies (both promoter and ORF) are scaled relative to it. Depicted are means + S.D. (N= 4; qPCR=8). Data are from Zhang et al, 2014 and are used with permission. **C.** ChIP-qPCR analysis of Pol II occupancy within the *YFR057w* promoter and ORF in *sir2Δ* and *SIR^+^* cells bearing the *ctk1Δ* deletion conducted as in **B**. Shown are means ± S.D. (N= 2 or 3; qPCR=4 or 6).

Is the severe reduction in heterochromatic transcription a consequence of impaired Pol II recruitment or its impaired release from the promoter? As shown in Figure 5C, Pol II occupancy of the heterochromatic *YRF057w* promoter and ORF was reduced to undetectable levels in the *ctk1Δ* mutant, and this was the case at all time points examined (*SIR^+^* panels). By contrast, Pol II levels were detectable at the heterochromatic *YFR057w* promoter and ORF in a *CTK1^+^* background, particularly at the longer time points (Figure 5B), consistent with the increase in *YFR057w* RNA levels. Moreover, Pol II occupancy of the euchromatic *YRF057w* gene in a *ctk1Δ* background was only 2- to 3-fold diminished (compare Figure 5C with 5B [*sir2Δ* panels]). Comparable analyses of *ctk1Δ bur1-as* (analogue sensitive) double mutants treated with the ATP analogue NM-PP1, whose Ser2 CTD phosphorylation is reduced to undetectable levels (Qiu et al., 2009), gave similar results (Figure 6, A and B). Importantly, cell viability was not impacted during the CX time course (Suppl. Fig. S1). Collectively, these experiments suggest that Ser2 phosphorylation of the Pol II CTD is critical to the transcription of the heterochromatic *YRF057w* gene and this PTM is important for Pol II recruitment and/or retention at this gene.

**Figure 6.**
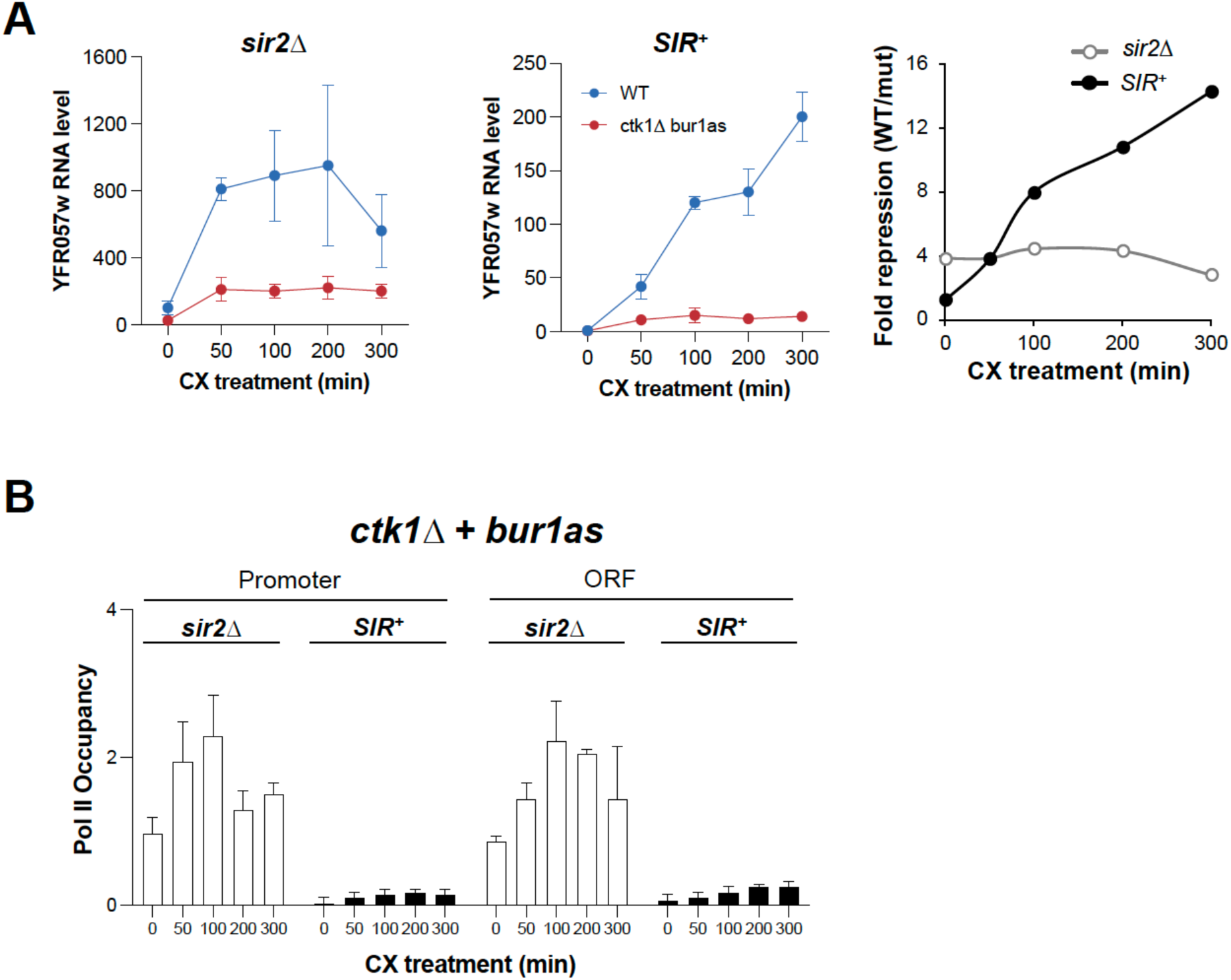
Heterochromatic gene activation is nearly abolished in cells deficient for CTD Ser2 phosphorylation. **A.** Expression assays were conducted as in Figure 5A except WT (*CTK1 BUR1*) *sir2Δ* and *SIR^+^* cells or isogenic *ctk1Δ bur1-as* double mutants were employed (strains HZY103, HQY1223, HZY203 and HQY1220, respectively). Following 1 h pretreatment with 15 μM 1-NM-PP1, cells were exposed to 200 μg/ml cycloheximide for the indicated times and then RNA was extracted. Depicted are means ± S.D. (N= 3; qPCR=6). **B.** ChIP-qPCR analysis of Pol II occupancy in *sir2Δ* and *SIR^+^* cells bearing the *ctk1Δ bur1-as* double mutation. Cells were treated as in **A**; Pol II ChIP analysis was conducted as in Figure 5C (N= 3; qPCR=6).

## Discussion

### An alternative pathway in trans-activation of the subtelomeric *YFR057w* gene

Previous work demonstrated that in a heterochromatic context, transcription can take place in the virtual absence of chromatin modifications typically linked to transcriptional activation (Zhang et al., 2014). These prior observations suggested that heterochromatic gene activation may rely on one or more alternative pathways. Here we have demonstrated that heterochromatic gene activation, as epitomized by the transgenic *hsp82-2001* gene, occurs in >50% of cells in the population. Moreover, this is likely the consequence of frequent transcription given that the level of expression in single cells is quite high.

One potential mechanism contributing to heterochromatic gene activation was tested here, namely the phosphorylation state of the Pol II CTD. We observed substantial phosphorylation of Ser2, Ser5 and Ser7 residues within the Pol II CTD at the cycloheximide-induced *YFR057w* gene. The presence of Ser2-phosphorylated Pol II was particularly notable for two reasons. First, it was significantly enriched at the native heterochromatic gene compared to what was observed when the gene was placed in a euchromatic context. Second, ablation of Ctk1, the kinase responsible for the bulk of Ser2 phosphorylation (Qiu et al., 2009; Bataille et al., 2012), reduced *YRF057w* mRNA levels >90% and rendered Pol II occupancy undetectable. This contrasts with previous findings that deletion of *CTK1* has little effect on Pol II occupancy at euchromatic genes (Ahn et al., 2004; Bataille et al., 2012). However, we also observed a reduction in Pol II occupancy at euchromatic *YFR057w* in both *ctk1Δ* and *ctk1Δ + bur1-as* mutants, although not to the same degree as seen at the natural, *SIR*-regulated *YFR057w* gene. Therefore, *YFR057w* may have an unusually strong requirement for Ser2 CTD phosphorylation and this requirement might be heightened by its heterochromatic state. While the basis for this remains unclear, it is possible that it is linked to the enhanced ability of the Ser2-phosphorylated CTD to recruit Pol II elongation factors such as Spt6 and Paf1C (Yoh et al., 2008; Sun et al., 2010; Tomson and Arndt, 2013) that may be particularly required for Pol II to traverse through *SIR*-stabilized nucleosomes (Wang et al., 2015) located within the *YRF057w* transcribed region. A model summarizing the potential differences in the mechanism by which euchromatic and heterochromatic genes activate transcription is presented in Figure 7.

**Figure 7.**
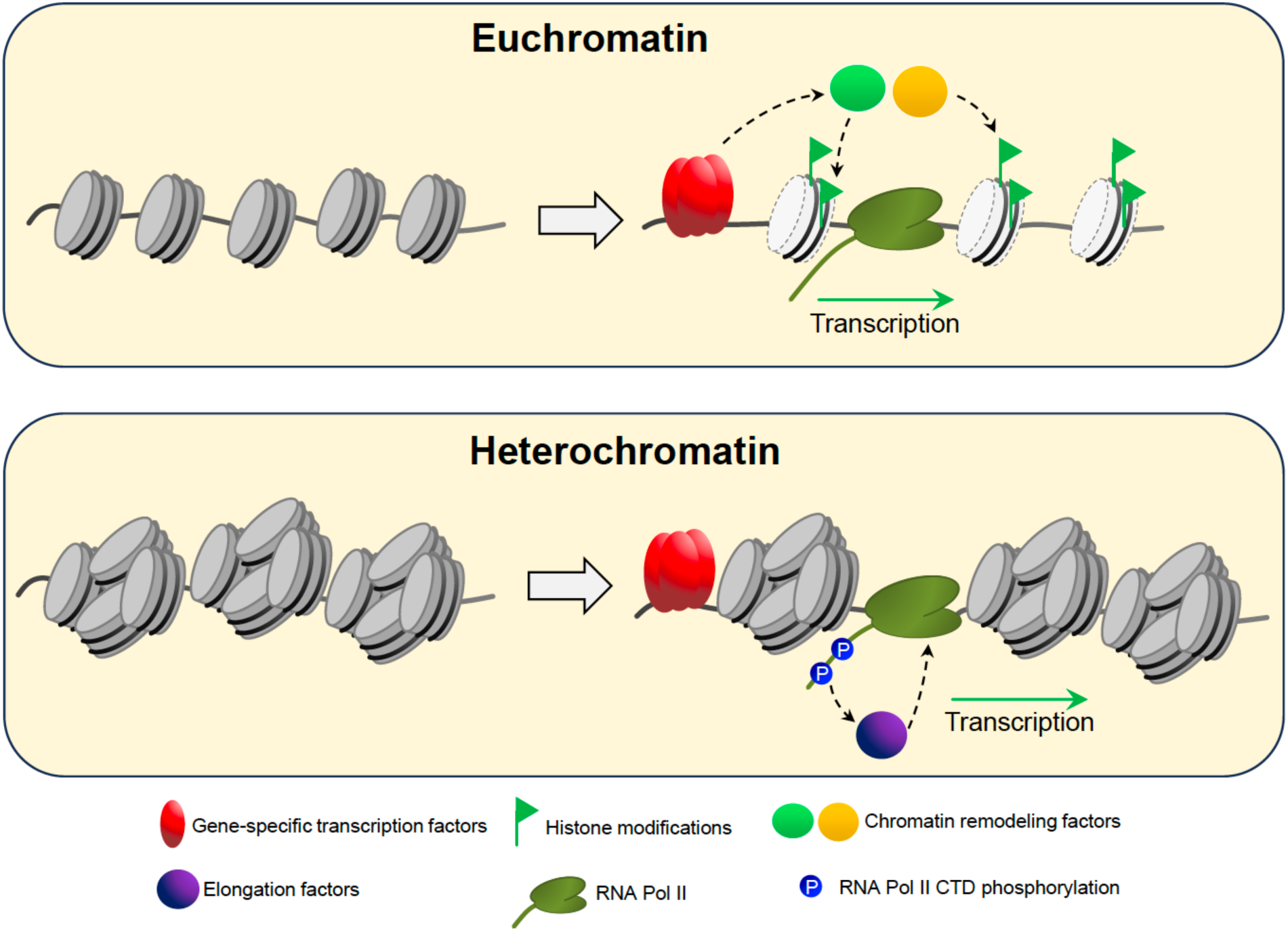
(*Top*) Gene-specific transcription factors activate transcription in euchromatin, facilitated by activating covalent histone modifications that serve as docking sites for transcriptional coactivators and chromatin remodeling complexes. (*Bottom*) Gene-specific transcription factors with sufficient activation potency override heterochromatin-mediated silencing and recruit key components of the transcription machinery. CTD phosphorylation of elongating Pol II then acts as an interacting moiety for elongation factors such as Paf1C and Spt6 that enable productive transcription.

To summarize, our previous results indicated that conventional histone modifications linked to gene expression are largely bypassed during heterochromatic gene activation. This is the case even though the same genes whose nucleosomal histones fail to be acetylated or methylated (at H3K4, K36 or K79) in a heterochromatic context are heavily modified by these same marks in a euchromatic context (Zhang et al., 2014). Alternative mechanisms must therefore exist to ensure the precise and dynamic regulation of Pol II genes embedded in heterochromatin, and here we have shown that at the subtelomeric *YFR057w* gene, Ser2 phosphorylation of the Pol II CTD is critically required. It is worth emphasizing that CTD phosphorylation does not by itself fully compensate for the absence of histone PTMs or other chromatin alterations that typically accompany euchromatic gene activation, since transcript levels of heterochromatin-assembled genes are consistently several-fold lower than their euchromatin-assembled counterparts. Moreover, additional activation pathways may be utilized at *YFR057w* and other heterochromatic genes. Indeed, the specific pathway(s) utilized by dynamically regulated heterochromatic genes – such as heat stress-inducible genes in *Arabidopsis* and actively transcribed heterochromatic genes in *Drosophila*, parasitic protozoa and mammals (reviewed in Zhang et al., 2014) – may prove to be promoter- and gene-specific.

## Materials & Methods

### Yeast Strains

The *hsp82* transgenic strains (SLY101 background) and isogenic WT, *ctk1Δ* and *ctk1Δ bur1-as* strains (BY4741 background) have been previously described (Sekinger and Gross, 1999; Qiu et al., 2009). Strain ASK101 was made by inserting the *NLS-eGFP-LEU2* cassette at the euchromatic *hsp82-2001* locus (replacing the ORF) in strain EAS2001 using homologous recombination. This resulted in the replacement of 2130 bp of *HSP82* with 3474 bp of the cassette. To obtain strain A6A, ASK101 with episomal expression of *SIR4* was crossed with LG1101. The resultant diploid was sporulated, tetrads were dissected and progeny with a Leu^+^ His^-^ genotype was selected and named A6A. Strains HQY1190, HQY1220 and HQY1223 were generously provided by J. Dong and A.G. Hinnebusch (NIH). Strains used in this study are listed in Table 1.

**Table 1.**
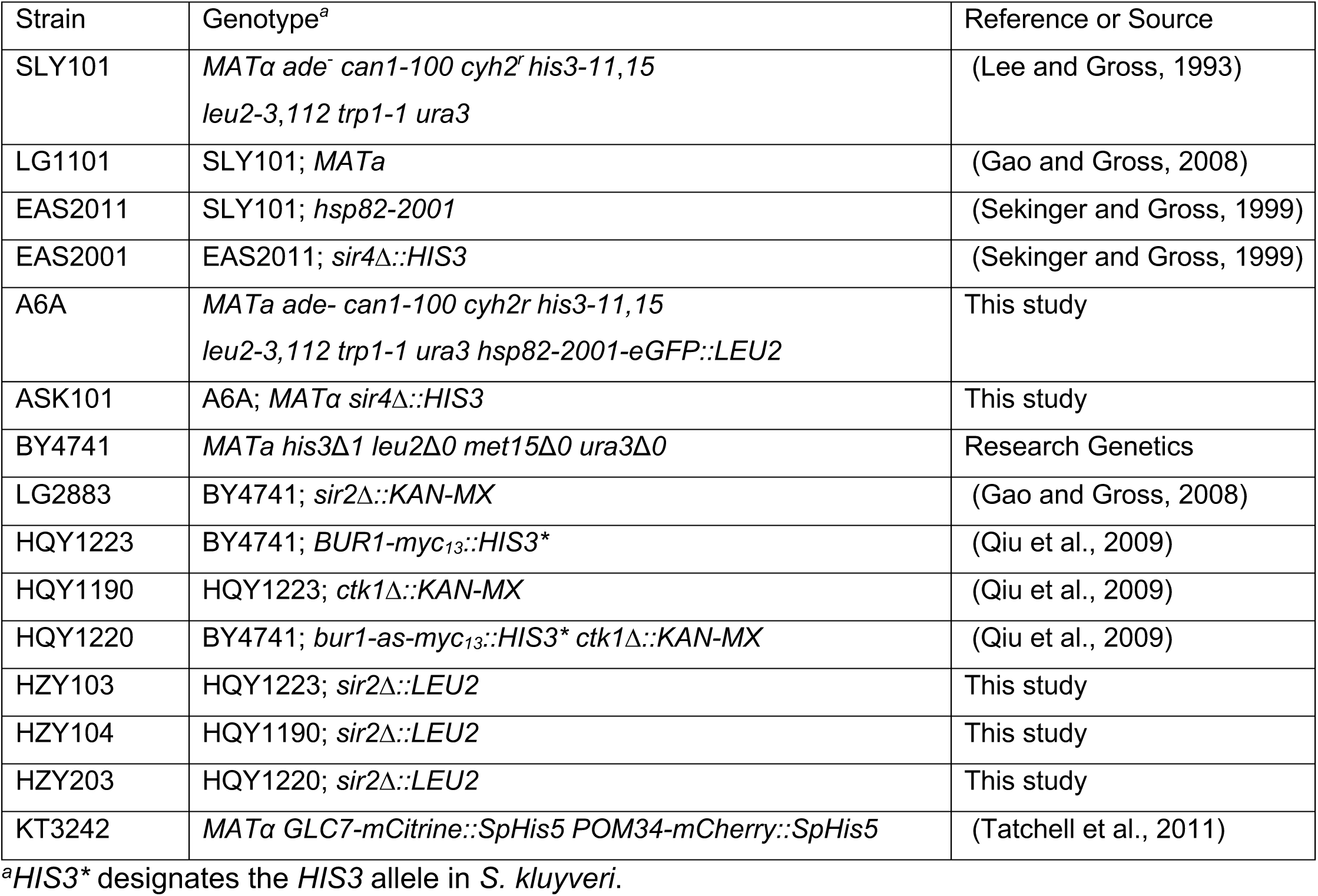
Yeast Strains.

### Cultivation Conditions

For experiments using the *ctk1Δ* strain HQY1190 and its isogenic wild-type counterpart HQY1223, cells were cultivated to early log phase (A_600_ ∼0.6) in rich yeast extract-peptone-glucose broth supplemented with 0.03 mg/ml adenine (YPDA) medium. Cycloheximide was then added to a final concentration of 200 μg/ml as previously described (Zhang et al., 2014). For experiments using the *ctk1Δ bur1-as* strain HQY1220 and its isogenic wild-type counterpart, HQY1223, cells were cultivated as usual, then 1-NM-PP1 (4-amino-1-tert-butyl-3-(1’-naphthylmethyl)pyrazolo[3-4-d]pyrimidine) (Toronto Research Chemicals, Inc.; no. A603003) was added to a final concentration of 15 μM, and cells were incubated for 1 hr at 30°C. At that point, a 10 ml aliquot was removed and 200 μg/ml CX was added to the remaining NM-PP1-treated cells. At 50, 100, 200 and 300 min, additional 10 ml aliquots were removed and processed as above.

### Fluorescence Microscopy

For cell imaging, cells were grown at 30°C to early log phase in YPDA, subjected to instantaneous heat shock at 39°C for the indicated times, and then fixed in 1% formaldehyde for 10 min. Following washing with phosphate-buffered saline (PBS) (pH 7.4), a small quantity was transferred to a patch of 2% agarose (prepared in PBS) on a glass slide. Cells were imaged on an Olympus fluorescence microscope with a UPlanFl 100 /1.3-numerical-aperature (NA) objective using a CoolSnap HQ charge-coupled-device camera. For imaging GFP, a 41001 filter set was used (Chroma Technology). To control camera acquisition and the Z axis stepping motor (Ludl Electronic Products), we used Slidebook version 4 software (Intelligent Imaging Innovations). Fluorescence images (binned 2 by 2) were acquired in a single plane. Background signal was calculated from the fluorescence levels of field view outside the cells. Cells with signal above the background were deemed fluorescence positive.

### Flow Cytometry Analysis

*S. cerevisiae* cells were grown at 30°C in YPDA to early log phase. Half of the 100 ml culture was subjected to an instantaneous 30°C to 39°C heat shock by mixing an equal volume of culture with pre-warmed medium (50°C) in 39°C water bath; the other half was maintained at 30°C (NHS control). For the HS sample, following 60 min incubation at 39°C, cells were transferred to 30°C for 30 min to allow folding and maturation of eGFP. Formaldehyde was then added to both HS and NHS cultures to a final concentration of 1% for 10 min. Cells were then collected from 1.5 ml of culture by centrifugation. The pellet was washed twice with 1 ml of PBS, and cells were resuspended in 1 ml PBS. The sample was sonicated briefly (15 sec at 10% amplitude using a Branson 250 Sonifier equipped with a microtip) to homogenize the suspension. Flow cytometry was conducted using a BD Biosciences LSRII-SORP Flow Cytometer (log forward scatter (FSC) 260V; SSC 119V). The eGFP fluorochrome was excited at 488nm and emission was collected through 530/30 nm bandpass filter. BD Biosciences FacsDiva software was used for acquisition and Floreada.io (https://floreada.io/) was used for making 2D spread plots. To calculate the change in expression of eGFP between NHS, 20 min and 60 min HS samples, the Overton cumulative histogram subtraction algorithm (Overton, 1988) was used in the FCS Express software suite.

### ChIP-qPCR

ChIP was conducted essentially as described (Zhang et al., 2014). 125 ml of mid-log cell culture were used in time course experiments, and 25 ml aliquots were removed for each time point. Two ml of soluble crosslinked chromatin were obtained from each cell sample, and ∼10% of that (200-250 µl) was employed for each IP. For phospho-CTD ChIP analysis, chromatin isolation and subsequent IP were carried out in the presence of phosphatase inhibitors (5 mM NaN_3_, 5 mM NaF and 2 mM Na_3_VO_4_) (Mayer et al., 2010).

Immunoprecipitations were conducted through addition of 40 µl of a 50% slurry of CL-4B Protein A Sepharose beads, with incubation at 4°C overnight. Exceptions to this were the phospho-CTD IPs using rat monoclonal Abs (see below), which were conducted in the presence of Protein G Sepharose 4 beads (GE Healthcare). For phospho-CTD ChIP, total Pol II (Rpb1) and phospho-CTD levels were both normalized to their respective levels at *PMA1* prior to calculating phospho-CTD/Rpb1 quotients. The following antibodies were used: Pol II, rabbit antiserum raised against an *E. coli* expressed GST-mouse Pol II CTD fusion protein with 52 heptad repeats (Zhao et al., 2005); Ser2-P CTD, Ser5-P CTD and Ser7-P CTD (rat monoclonal Abs 3E10, 3E8, and 4E12, respectively; Helmholtz Zentrum, Munich). Amplicons used for Real Time PCR (coordinates relative to ATG) were as follows: *YFR057w* promoter, -115 to -45; *YFR057w* ORF, +312 to +437; *PMA1* 5’ coding region, +49 to +112.

### RNA assays

RT-qPCR was conducted as previously described (Zhang et al., 2014). The Pol III transcript *SCR1* was used to normalize *YFR057w* mRNA levels. Primers were designed to target the ORF of *YFR057w* (+312 to +437) and the body of *SCR1* (+385 to +483).

## Acknowledgements

We thank Dr. Aseem Z. Ansari (St. Jude Children’s Research Hospital) for helpful discussions and Drs. Jinsheng Dong and Alan G. Hinnebusch (NIH) for the gift of yeast strains. This work was supported by grants from the NSF (MCB-1158516) and NIH (R01 GM138988) awarded to D.S.G.

## Figure Legends

**Suppl. Figure S1.** Cycloheximide 1-NM-PP1 viability assay.

One portion of each *SIR2* and *sir2Δ* culture (BY4741 background) was subjected to a 1 h pretreatment with 15 μM 1-NM-PP1 (blue bars), the other was not (yellow bars). A_600_ readings were taken either prior to (red bars) or 300 min following addition of 200 μg/ml cycloheximide (yellow and blue bars). Mean values are shown; N= 2 independent cultures.

