## Supplemental Figures for "A critical role for Pol II CTD phosphorylation in heterochromatic gene activation"

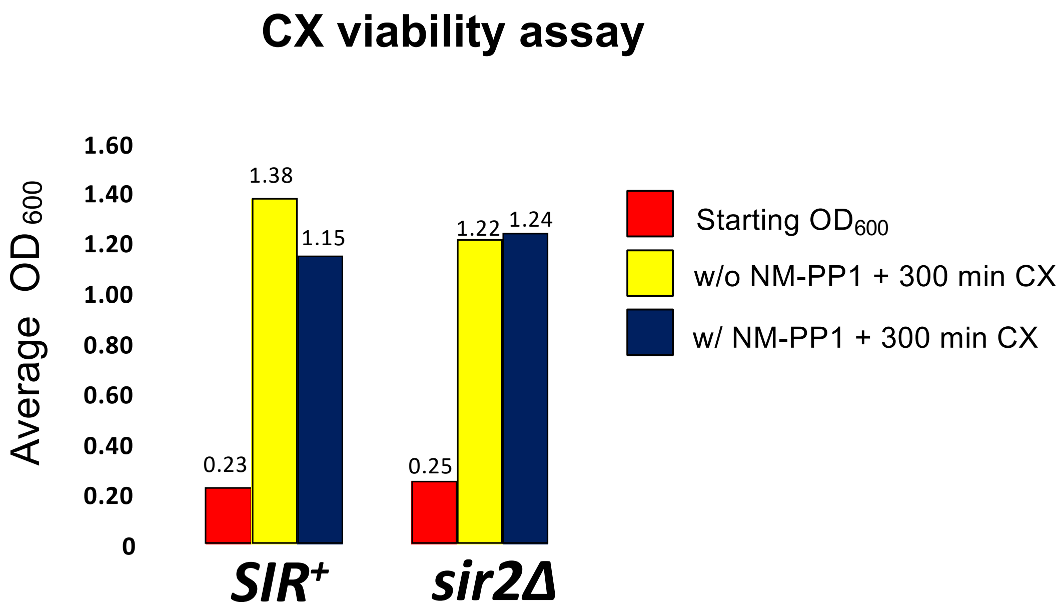


**Suppl. Figure S1. Cycloheximide 1-NM-PP1 viability assay.**

One portion of each *SIR*$^+$ and *sir2*Δ culture (BY4741 background) was subjected to a 1 h pretreatment with 15 μM 1-NM-PP1 (blue bars), the other was not (yellow bars). $A_{600}$ readings were taken either prior to (red bars) or 300 min following addition of 200 μg/ml cycloheximide (yellow and blue bars). Mean values are shown; N= 2 independent cultures.
